# Integrative Analysis of Single-cell and Bulk RNA Sequencing Reveals Macrophage Heterogeneity in Lyme Arthritis

**DOI:** 10.1101/2025.02.14.638349

**Authors:** Yan Dong, Yantong Chen, Yanshuang Luo, Meng Liu, Yingchen Zhou, Yutian Yang, Xuesong Chen, Xiaorong Liu, Fusong Yang, Zhi Liang, Liye Zi, Hongbing Gao, Zhichao Cao, Fucun Gao, Chao Song, Guozhong Zhou

**Author notes:** Correspondence to:* Chao Song, Department of Pain Medicine, The Affiliated Anning First People’s Hospital of Kunming University of Science and Technology, Kunming, Yunnan Province, 650302, China, Guozhong Zhou, Department of Pain Medicine, The Affiliated Anning First People’s Hospital of Kunming University of Science and Technology, Kunming, Yunnan Province, 650302, China; School of Basic Medical Sciences, Kunming University of Science and Technology, Kunming, Yunnan Province, 650500, China. Yan Dong and Yantong Chen contributed equally to this work.

## Abstract

**Introduction:** Lyme arthritis (LA) is a common late-stage manifestation of Lyme disease caused by *Borrelia burgdorferi* (Bb) infection. Macrophages play a crucial role in LA pathogenesis, yet their heterogeneity and functional states within the arthritic microenvironment remain poorly understood.

**Methods:** We integrated single-cell RNA sequencing data from ankle joint tissues of Bb-infected C57BL/6 mice with bulk RNA sequencing data from both infected ankle joints and bone marrow-derived macrophages. We developed a LA mouse model and validated key findings through histopathological examination and RT-qPCR analysis.

**Results:** We identified nine distinct cell types in the joint microenvironment with significant compositional shifts during disease progression. Analysis of 16,617 Macrophages/Monocytes revealed seven functionally distinct subpopulations with differential responses to infection. Pseudo-time trajectory analysis demonstrated three principal differentiation pathways from monocyte-derived macrophages toward anti-inflammatory, TNF+, and IL1b+ inflammatory phenotypes. Cell-cell communication analysis revealed significantly altered TGF-β signaling networks in LA, with the Tgfb1-(Tglbr1+TgIbr2) axis playing a critical role. GSVA and pathway analyses showed metabolic reprogramming from oxidative phosphorylation toward glycolysis in macrophages during infection. Through integrated analysis and LASSO regression, we identified four characteristic genes (SIRPB1C, FABP5, MMP14, and EGR1) as potential LA biomarkers and developed a diagnostic nomogram with high predictive accuracy.

**Conclusion:** Our integrative analysis provides comprehensive insights into macrophage heterogeneity and functional plasticity in LA, identifying characteristic biomarkers and potential therapeutic targets. The metabolic reprogramming and altered TGF-β signaling networks we identified may contribute to disease pathogenesis and offer new avenues for intervention strategies.

## 1. Introduction

Lyme arthritis (LA), the most common and persistent late-stage manifestation of Lyme disease, develops from infection with the tick-borne spirochete Borrelia burgdorferi (Bb), typically emerging months after initial infection[1,2]. Characterized by intermittent or persistent swelling and pain predominantly in large joints, especially the knee, LA significantly impairs quality of life. Without antibiotic treatment, approximately 60% of Lyme disease patients develop arthritis[3,4]. The condition’s complexity arises from pathogen-host immune system interactions that trigger inflammation and, in antibiotic-resistant cases, chronic arthritis[5,6]. Despite extensive research into immune responses, bacterial components, and genetic susceptibility, the complete pathophysiological mechanisms of LA remain incompletely understood.

Extensive research has shown that macrophages play a central role in the pathogenesis of LA[7,8]. Upon Bb infection, they secrete CCL4 and CCL2 chemokines that recruit monocytes and T cells to infection sites while amplifying inflammation through phagocytosis and antigen presentation[9,10]. Both animal and human studies confirm that macrophages induce arthritis directly and through pro-inflammatory cytokines (IL-1β, TNF-α, IFN-γ), which maintain and exacerbate joint inflammation[11–13]. However, macrophages also contribute to *Borrelia* clearance and inflammation resolution[14]. Their activity is regulated by polarization states, cytokine environment, and immune cell networks, making a comprehensive understanding of their role in LA pathogenesis essential.

With the rapid development of single-cell RNA sequencing (scRNA-seq) technology, researchers have been able to analyze tissue heterogeneity at the single-cell level[15–17], facilitating precise studies on the roles of individual cells such as macrophages in the progression of LA. However, studies integrating scRNA-seq with bulk RNA sequencing to investigate macrophage roles in LA remain scarce. In this study, we established a Bb-infected C57BL/6 mouse model and performed transcriptome sequencing of ankle joint tissues to analyze gene expression changes during arthritis development. We implemented a multi-omics integrative approach, correlating mouse joint tissue transcriptome data with publicly available bone marrow-derived macrophage transcriptome and scRNA-seq datasets to construct a comprehensive gene expression network elucidating host immune responses to Bb infection. This multidimensional approach enhances our understanding of macrophages’ central role in LA pathogenesis and provides theoretical foundation for developing targeted therapies.

## 2. Methods

### 2.1 Single-cell RNA Sequencing Data Quality Control and Processing

The scRNA-seq data of LA was accessed from the GSE233850 dataset, which is established in the GEO database corresponding to a published article “Single-cell RNA sequencing of murine ankle joints over time reveals distinct transcriptional changes following *Borrelia burgdorferi* infection” [18]. The single-cell RNA sequencing data processing workflow is as follows: Raw data was obtained from ankle joint samples of 1 normal and 4 *Borrelia burgdorferi*-infected C57BL/6J mice from the GSE233850 dataset. Quality control criteria include: more than 500 expressed genes per cell, each gene expressed in at least 3 cells, and mitochondrial RNA content less than 20%[19]. Data processing using the Seurat package: Normalization was performed using the NormalizeData function (natural logarithm of expression values multiplied by total gene expression ×10000), followed by identification of 2000 highly variable genes using FindVariableFeatures, data centering using ScaleData, and finally data integration using FindIntegrationAnchors (RunHarmony) with 2000 anchors to eliminate batch effects. A total of 42,385 cells were obtained for subsequent analysis[20].

### 2.2 Dimensionality Reduction and Data Clustering

To visualize high-dimensional gene expression data effectively[21,22], we implemented a sequential dimensionality reduction and clustering protocol. We first performed principal component analysis on highly variable genes using RunPCA, followed by data integration with RunHarmony using 2000 anchor points. We then applied FindNeighbors and FindClusters functions for clustering analysis. Finally, we employed RunUMAP (dims=1:14, resolution=0.5) for dimensionality reduction visualization to display cell clustering results.

### 2.3 DEG, Marker Gene and Active Subgroups Identification

To characterize each cluster, a comprehensive manual annotation was performed. Using the Seurat package’s FindAllMarkers function, we identified differentially expressed genes (DEGs) with |log2FoldChange|>1 and adjusted P-value<0.05, and determined cell type-specific marker genes. We employed the AUCell R package to calculate pathway activity scores in single-cell data[23]. After extracting the RNA expression matrix from our Seurat object with GetAssayData, we ran AUCell_run with predefined gene sets to quantify pathway enrichment per cell. For pathways of interest, we extracted area under the curve (AUC) values and incorporated them into the Seurat metadata.

### 2.4 GO and KEGG Pathway Enrichment Analysis

We conducted GO functional enrichment and KEGG pathway enrichment analyses on differentially expressed genes using the clusterProfiler package (v3.14.3) and visualized results with the ggplot2 package[24,25].

### 2.5 Pseudo-time Analysis

To analyze the continuous dynamic changes in cell types and associated gene expression, we used Monocle 3 to reconstruct cellular developmental trajectories from single-cell data[26]. We extracted six macrophage subtypes (Tnf+ inflammatory, anti-inflammatory, monocyte-derived, Il1b+ inflammatory, tissue-repair, and monocyte-derived macrophages) to create a cell_data_set object. Following PCA preprocessing, we removed batch effects with the align_cds function using orig.ident as the alignment group. We performed dimensionality reduction via reduce_dimension and applied low-resolution clustering to ensure distinct trajectories per cluster. We constructed the cellular trajectory network using learn_graph and employed a custom function to automatically identify monocyte-derived macrophages as developmental origin points. We then ordered cells along the developmental trajectory with order_cells and visualized both pseudo-time progression and expression patterns of the top five genes using plot_cells and plot_genes_in_pseudotime functions.

### 2.6 Cell-Cell Communication Analysis

Using the CellChat package (v1.0.0) to analyze intercellular communication networks, cell communication relationships are constructed based on expression data of known ligand-receptor pairs to infer interaction patterns and their abundance between different cell types[27].

### 2.7 C57BL/6 Mouse LA Modeling and Transcriptome Sequencing

Mice: Female C57BL/6J mice (22-24g, 4-6 weeks old) from Beijing Vital River Laboratory Animal Technology Co., Ltd. were housed in SPF-grade facilities at Kunming Medical University’s School of Basic Medicine with 12-hour light/dark cycles at 21.5°C, free food and water access, and daily bedding changes. The study was approved by Anning First People’s Hospital Medical Ethics Committee (Approval No.2024-046-01). Bacteria propagation and infection: *Borrelia burgdorferi* strain B31 (ATCC-35210) was cultured in complete BSK-II medium at 37 °C with 5 % CO_2_ and 100 % humidity. Experimental mice received bilateral footpad injections of spirochete suspension (1×10^6/50μl per side), while controls received equivalent PBS volumes. Ankle joint swelling was measured biweekly using calipers, with X-ray examinations performed on randomly selected mice. Sample Collection and Transcriptome Analysis: On day 16 post-surgery, we sampled 6 mice per group. Half the samples underwent histopathological examination with 4 μm paraffin sections for HE staining. The remaining samples (approximately 80 mg of ankle joint tissue) were sent to LC Bio Technology CO., Ltd. (Hangzhou, China) for transcriptome sequencing using the Illumina Novaseq 6000 platform with paired-end sequencing (PE150 mode). Bioinformatics analysis was conducted on the standardized clean data.

### 2.8 Transcriptome Data Integration

Based on our research focus on the role of macrophages in LA synovial tissue, we obtained transcriptome sequencing dataset (GSE125503, platform GPL17021) of Bb-infected macrophages from the Gene Expression Omnibus (GEO) database. This dataset includes 14 samples derived from bone marrow-derived macrophages (BMDMs) of C57BL/6 mice, comprising 4 control samples and 10 Bb-infected samples. Subsequently, we integrated our team’s Bulk-transcriptome data with the GSE125503 Bulk-transcriptome data. We merged different datasets by columns using the cbind function and obtained common genes through Reduce(intersect, geneList). Batch effects were then corrected using the ComBat function. Following this, we established criteria for identifying DEGs using the R package DESeq2. Specifically, we applied a threshold of |log_2_ FoldChange|> 1 and adjusted P-value <0.05 to select DEGs for subsequent analysis[28,29].

### 2.9 Least Absolute Shrinkage and Selection Operator (LASSO) Regression Analysis

We identified intersection genes between macrophage-related single-cell sequencing DEGs and our merged transcriptome dataset DEGs. Using these common differential genes, we constructed and validated a model using the Least Absolute Shrinkage and Selection Operator (LASSO) method[30].

### 2.10 Quantitative Reverse Transcription PCR (qRT-PCR)

Cell Culture and Treatment: RAW264.7 cells were purchased from Shanghai Zhongqiao Xinzhou Biotechnology Co., Ltd. The cells were infected with Bb (MOI=1) for 6 hours, or treated with vehicle control (PBS). Samples were stored at −80°C until analysis. RNA Extraction and qRT-PCR Analysis: Total RNA was extracted using MolPure^®^ Cell/Tissue Total RNA Kit (#19221ES50, ShangHai Yeasen Biotechnology) and reverse transcribed into complementary DNA using Hifair^®^III 1st Strand cDNA Synthesis SuperMix for qPCR (#11141ES60, ShangHai Yeasen Biotechnology). qRT-PCR was performed on the Gentier 96R Real-Time PCR System (Tian Long) using Hieff unicon universal blue Qpcr SYBR Master mix (#11184ES08, ShangHai Yeasen Biotechnology) under the following conditions: 95°C for 2 minutes, followed by 40 cycles of 95°C for 1 second and 60°C for 30 seconds. Expression of target genes was normalized to Gapdh using the 2^^(-ΔΔCT)^ method. The primers were synthesized by Tsingke Biotechnology (Beijing). The primer sequences (5’-3’) used to amplify SIRPB1C were forward, TCAGCAGACAGTGGCCTTTA and reverse, CTGAGAGCGGACATCCCTAG. The primers used to amplify FABP5 were forward, CGGGAAGGAGAGCACGATAA and reverse, TTGTTGCATTTGACCGCTCA. The primers used to amplify MMP14 were forward, AGGCCAATGTTCGGAGGAAG and reverse, GTGGCACTCTCCCATACTCG. The primers used to amplify EGR1 were forward, AGTGATGAACGCAAGAGGCA and reverse TAGCCACTGGGGATGGGTAA. The primers used to amplify GAPDH were forward, TGTTGCCATCAATGACCCCT and reverse TCGCCCCACTTGATTTTGGA.

### 2.11 Construction and Testing of LA Diagnostic Nomogram Model

Using the “pROC” and “rms” R packages[31,32], we developed a nomogram for LA diagnosis. Risk scores were calculated based on the expression levels of each candidate optimal feature gene, with the total risk score defined as the sum of all individual gene risk scores. The diagnostic value of this nomogram for LA was evaluated through decision trees, calibration curves, and ROC curves.

### 2.12 Statistical Analysis

Analyses and visualizations were performed using R (version 4.2.1) and GraphPad Prism (version 9.0.2). Data are presented as mean ± SEM. We compared groups using two-tailed Student’s t-test or Mann-Whitney U test, with significance defined as *P* < 0.05 (*), *P* < 0.01 (**), and ns for not significant.

**Figure 1.**
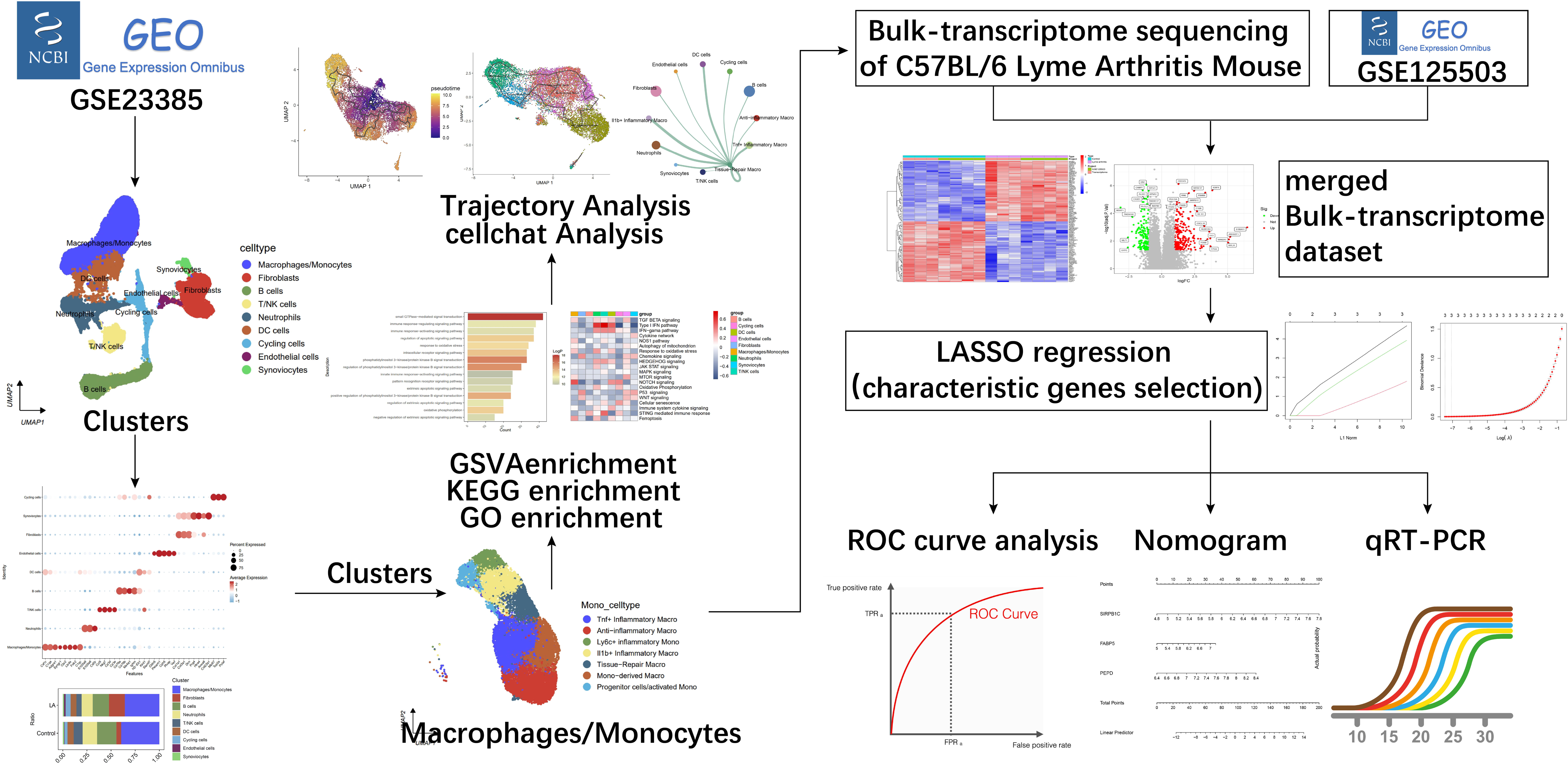
Comprehensive workflow diagram of this investigation.

## 3. Results

### 3.1 Single-cell Transcriptome of LA

We analyzed 5 samples from the GSE233850 dataset, including ankle joint samples from 1 uninfected normal C57 mouse and 4 paired Bb-infected C57 mice. After removing low-quality data, 42,385 cells were retained for analysis. Following normalization and PCA, we retained the first 14 dimensions and visualized clustering using UMAP, dividing cells into 20 clusters (Figure 2A). SingleR package analysis identified 9 distinct cell types (Figure 2B). Figures 2C and 2D showed the single-cell transcriptome atlas of normal and LA samples and across samples, respectively. The cell population comprised: Macrophages/Monocytes (16,617 cells, 39.2%; high expression of Csf1r, C1qa, Adgre1, Syngr1), Neutrophils (4,068 cells, 9.6%; S100a9, S100a8, Csf3r), T/NK Cells (2,472 cells, 5.8%; Ccl5, Nkg7, Cd3d, Cd3e), B Cells (7,582 cells, 17.9%; Cd79a, Cd79b, Ms4a1, Ighm), DC Cells (2,492 cells, 5.9%; H2-Eb1, Klrd1, Slamf7), Endothelial Cells (608 cells, 1.43%; Cldn5, Pecam1, Cdh5), Fibroblasts (5,999 cells, 14.2%; Col1a1, Col3a1, Dcn), Synoviocytes (475 cells, 1.1%; Prg4, Htra4, Anxa8), and Cycling Cells (2,072 cells, 4.9%; Mki67, Top2a, Pclaf) as shown in Figures 2E-2G. Figure 2H showed their proportions between groups, with the LA model exhibiting higher percentages of Fibroblasts, Cycling cells, and Endothelial cells, while Macrophages/Monocytes, B cells, T/NK cells, Neutrophils, DC cells, and Synoviocytes showed relatively lower proportions.

**Figure 2.**
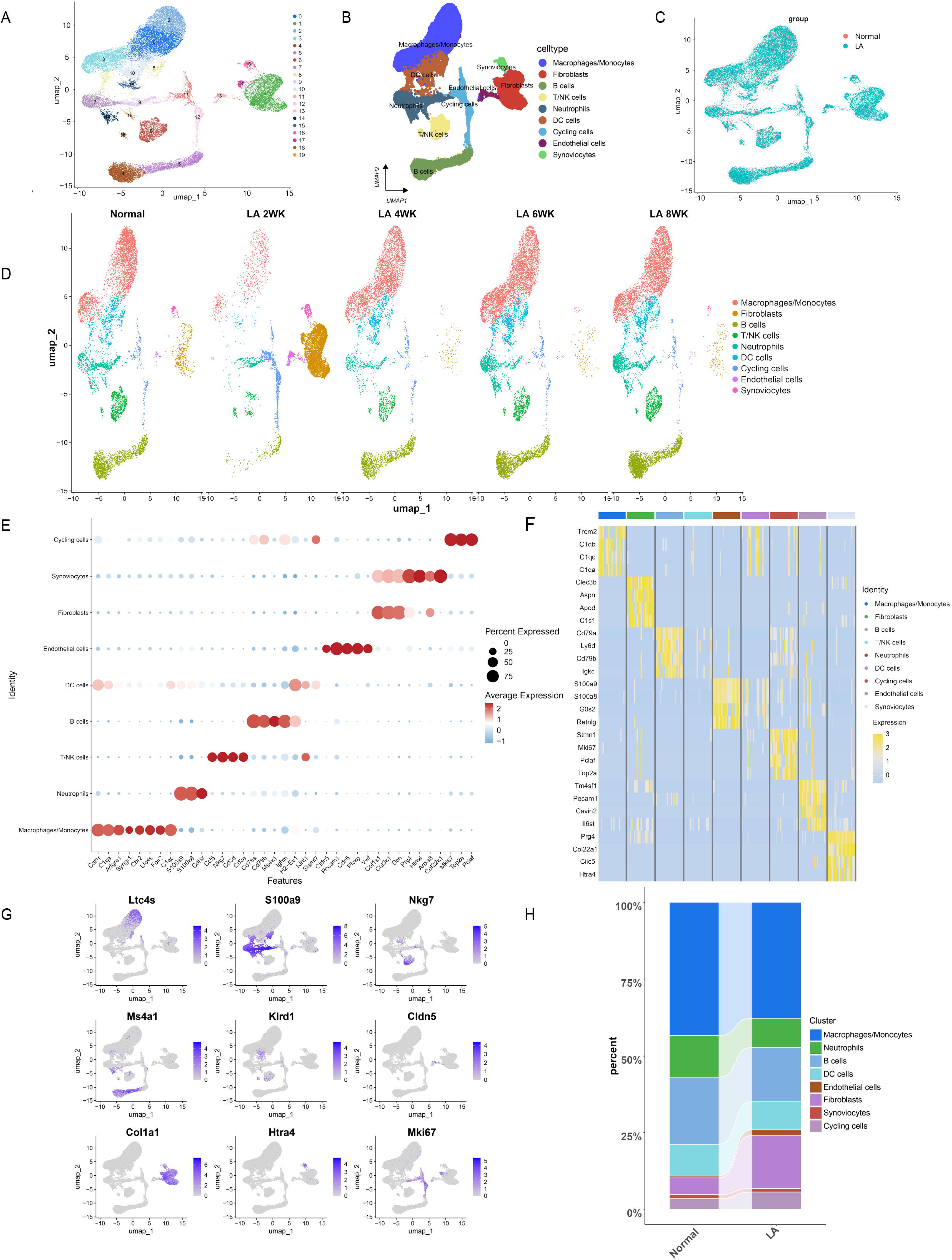
Single-cell atlas of normal and Lyme arthritis C57BL/6 mouse samples. (A) UMAP clustering of single cells into 20 clusters; (B) Nine cell types identified by marker gene expression; (C) UMAP comparison between normal and Lyme arthritis model groups; (D) UMAP distribution across 5 samples; (E) Bubble plot of marker gene expression in 9 cell types. Bubble plots display marker gene expression in cell subpopulations. Bubble color intensity (red) corresponds to expression level, while size reflects the number of expressing cells; (F) Heatmap of differential gene expression among cell types; (G) Histogram shows the cumulative distribution of selected marker genes of cell types; (H) Cell type proportion comparison between normal and Lyme arthritis groups.

### 3.2 Single-Cell GSVA Pathway Analysis of KEGG Metabolic Signatures

Figure 3 showed GSVA enrichment analysis using the AUCell algorithm, featuring differential gene expression across cell subpopulations (Figure 3A). We analyzed 20 KEGG signaling pathway activities within various cellular subsets (Figure 3B), with particular focus on glycolysis/gluconeogenesis and oxidative phosphorylation pathways visualized through violin plots (Figure 3C-D), UMAP projections (Figure 3E, 3G), and box plots (Figure 3F, 3H). Results showed increased glycolytic/gluconeogenic activity and decreased oxidative phosphorylation in Macrophages/Monocytes following Bb infection, indicating a metabolic shift from aerobic to anaerobic metabolism and highlighting metabolic reprogramming in host cells during pathogenic challenge.

**Figure 3.**
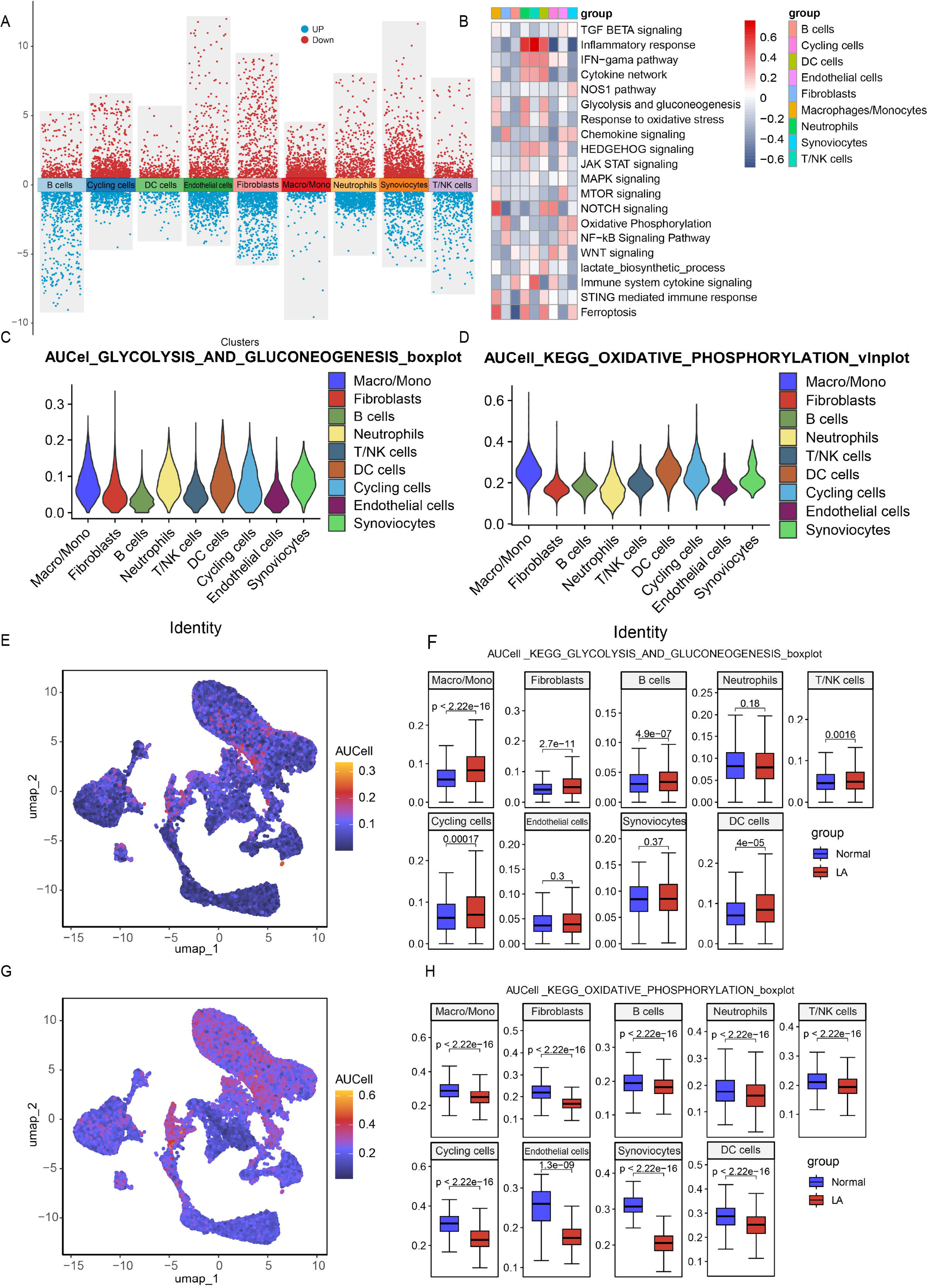
GSVA Enrichment Analysis Between Specific Subgroup Cells and Groups. (A) Volcano plot of differential gene expression across cellular subpopulations. Red dots indicate significantly upregulated cluster-specific genes; blue dots show significantly downregulated genes; (B) GSVA enrichment analysis among 9 cell types; (C) Violin plot of AUC scores for glycolysis and gluconeogenesis pathway-related gene sets; (D) Violin plot of AUC scores for oxidative phosphorylation pathway-related gene sets; (E) UMAP colorogram showing glycolysis and gluconeogenesis pathway activity, with brighter colors indicating higher activity; (F) Box plot of AUC scores for glycolysis and gluconeogenesis pathway-related gene sets; (G) UMAP colorogram showing oxidative phosphorylation pathway activity, with brighter colors indicating higher activity; (H) Box plot of AUC scores for oxidative phosphorylation pathway-related gene sets.

### 3.3 Single-cell Transcriptome Landscape

Due to Macrophages/Monocytes’ critical immunoregulatory role in LA, we analyzed 16,617 of these cells from all samples. Figure 4 showed scRNA-seq analysis identified 7 distinct subtypes: Anti-inflammatory macrophages (3,331 cells; high expression of Olfml3, Sparc, Vsig4), TNF+ Inflammatory macrophages (3,186 cells; Tnf, Cxcl2, Ccl7), Tissue-Repair macrophages (2,052 cells; Vegfa, Arg1, Adam8), Mono-derived macrophages (2,030 cells; Mrc1, Gas6), Il1b+ Inflammatory macrophages (1,924 cells; Il1b, Clec4n, Ptgs2), Ly6c+ inflammatory Mono (1,034 cells; Ly6c2, Hp, Thbs1), and Progenitor cells/activated Mono (1,033 cells; Plac8, Slfn1). Comparison between normal and LA model groups (Figures 4C-4G) revealed significant compositional differences: the LA model showed decreased proportions of Anti-inflammatory macrophages, Mono-derived macrophages, and Ly6c+ inflammatory monocytes, while TNF+ Inflammatory macrophages, Il1b+ Inflammatory macrophages, Tissue-repair macrophages, and Progenitor cells/activated monocytes increased (Figure 4H).

**Figure 4.**
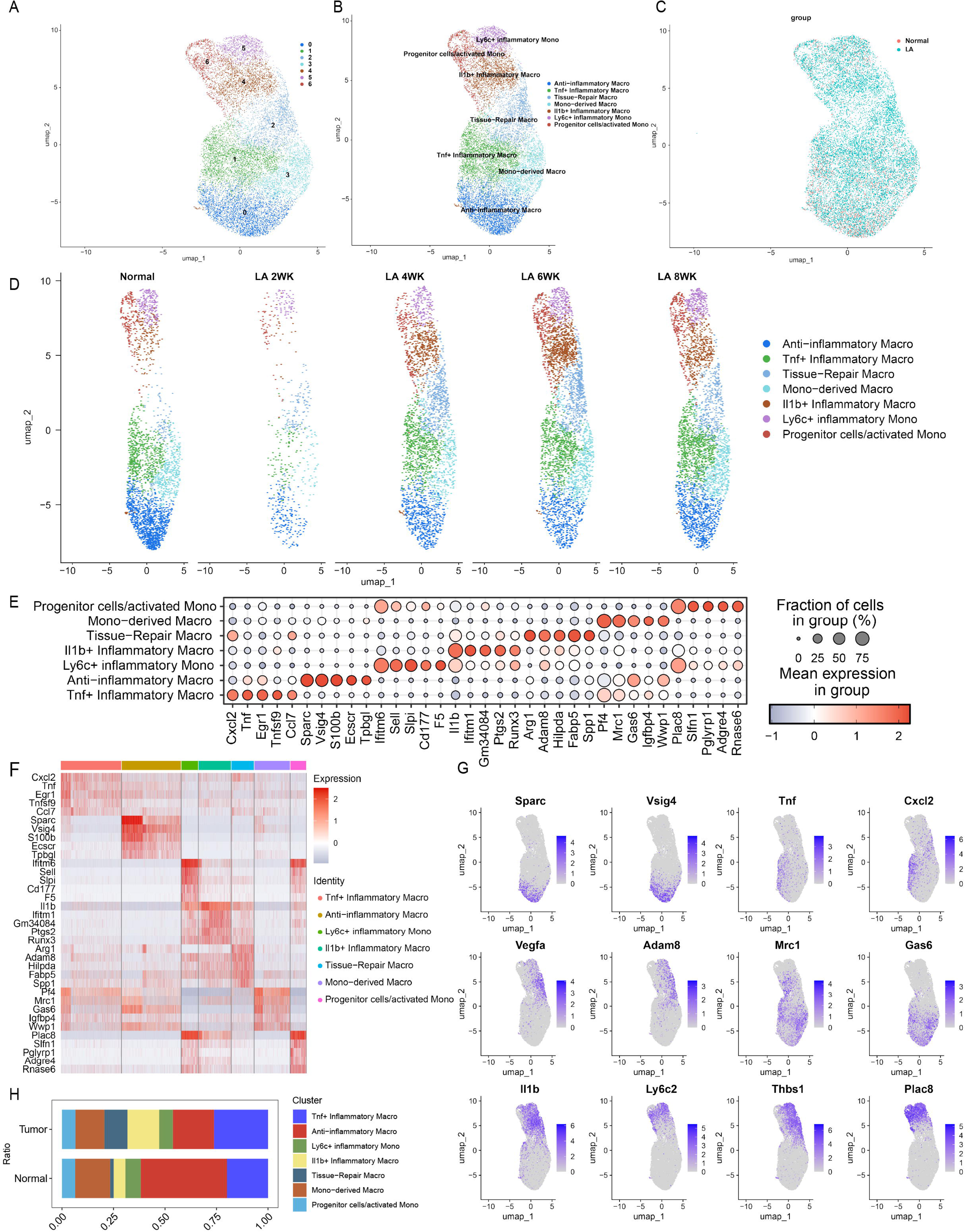
Single-cell transcriptome atlas of Macrophages/Monocytes. (A) UMAP plot showing 7 distinct Macrophage/Monocyte clusters; (B) UMAP plot with 7 Macrophage/Monocyte subtypes identified by signature molecular markers; (C) UMAP comparison between normal and Lyme arthritis model groups; (D) UMAP distribution across 5 samples; (E) Bubble plot showing differential expression patterns of signature genes across 7 cell subgroups; (F) Heatmap of differential gene expression among the 7 cell subgroups; (G) UMAP visualization of marker gene expression patterns for each cell subgroup; (H) Comparative proportions of cell subgroups between normal and Lyme arthritis groups.

### 3.4 Macrophage-Related Enrichment Analysis

To elucidate biological functions of Macrophage/Monocyte marker genes, we performed GO and KEGG enrichment analyses. GO functional enrichment of differentially expressed genes (Figure 5A-B) revealed: Biological Processes (BP) primarily related to endosomal transport, precursor metabolite and energy generation, hydrolase activity upregulation, cellular respiration, vesicle organization, and oxidative phosphorylation. Cellular Components (CC) mainly associated with mitochondrial protein complexes, vacuolar membranes, early endosomes, and lysosomal/endosomal structures. Molecular Functions (MF) included phospholipid binding, GTPase regulatory activity, nucleoside-triphosphatase regulation, active transmembrane transport, phosphatidylinositol binding, and GTPase interactions. KEGG pathway analysis (Figure 5C) showed signature genes enriched in inflammatory response pathways linked to energy metabolism regulation, including bacterial epithelial invasion, osteoclast differentiation, Toll-like receptor signaling, rheumatoid arthritis, carbon metabolism, HIF-1 signaling, glycolysis/gluconeogenesis, arginine/proline metabolism, and adherens junctions. Figure 5D shows differential analysis within Macrophages/Monocytes, while Figure 5E presents a diagonal volcano plot. GO enrichment bar plots for upregulated and downregulated genes (Figure 5F) revealed distinct patterns: upregulated genes showed significant enrichment in pathways governing precursor metabolite generation, nucleotide metabolism, phospholipid processes, and lipid/small molecule metabolic regulation. Downregulated genes showed predominant enrichment in oxidative phosphorylation, aerobic and cellular respiration, energy derivation through organic compound oxidation, and mitochondrial respiratory chain/NADH dehydrogenase complex assembly.

**Figure 5.**
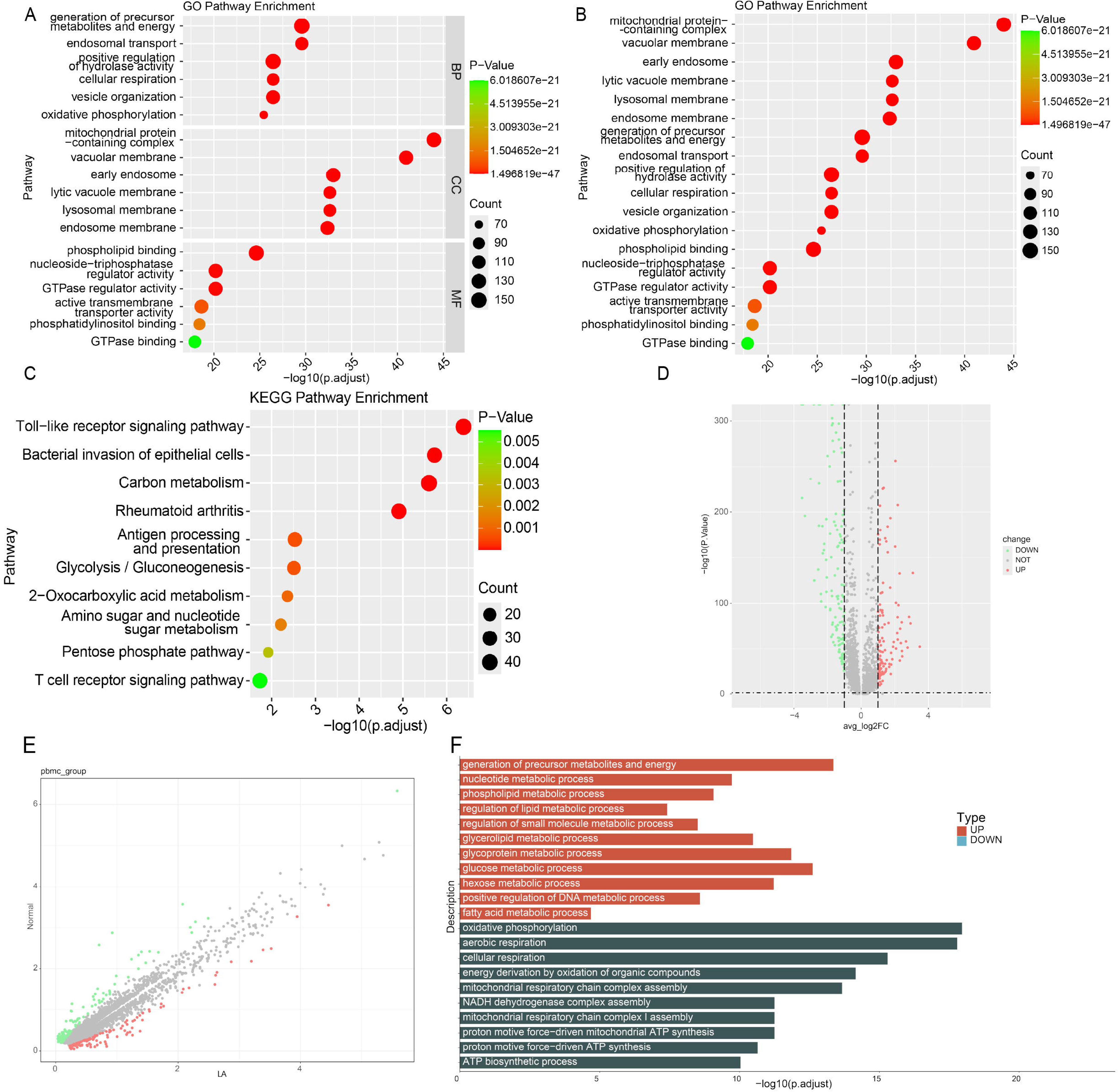
Enrichment Analysis Between Macrophage/Monocyte Subgroups. (A-B) GO functional enrichment analysis of differentially expressed genes in Macrophages/Monocytes, categorized by biological processes (BP), molecular functions (MF), and cellular components (CC); (C) KEGG pathway enrichment analysis showing functional distribution of differentially expressed genes across macrophage/monocyte populations; (D) Scatter plot comparing differential expression between Macrophage/Monocyte groups; (E) Diagonal volcano plots illustrating differential gene expression between Macrophage/Monocyte subpopulations; (F) Bar plots showing GO enrichment analysis of upregulated and downregulated genes across Macrophage/Monocyte subpopulations.

### 3.5 Pseudo-time Analysis

Using Monocle 3 R, we analyzed macrophage differentiation trajectories. Figure 6A showed dynamic RNA expression changes across five macrophage subgroups, while Figure 6B illustrateed pseudo-time progression from dark to light-colored cells. Our analysis identified monocyte-derived macrophages as the primary origin point, diverging into three distinct developmental paths: toward anti-inflammatory macrophages, Tnf+ inflammatory macrophages, and Il1b+ inflammatory macrophages. The trajectory map highlighted macrophage plasticity during disease progression, with tissue-repair macrophages primarily functioning as an intermediate state evolving toward Tnf+ inflammatory phenotypes rather than maintaining reparative functions (Figure 6C-6D). Figure 6E showed pseudo-time dynamics of C1qa, C1qb, and C1qc genes during LA-related macrophage development. These genes exhibit similar expression patterns: high expression in early pseudo-time (0-5), particularly in mono-derived, anti-inflammatory, and TNF+ inflammatory macrophages, followed by gradual decline in mid-to-late pseudo-time (5-10), with tissue-repair macrophages showing the most pronounced decrease and Il1b+ inflammatory macrophages maintaining intermediate expression.

**Figure 6.**
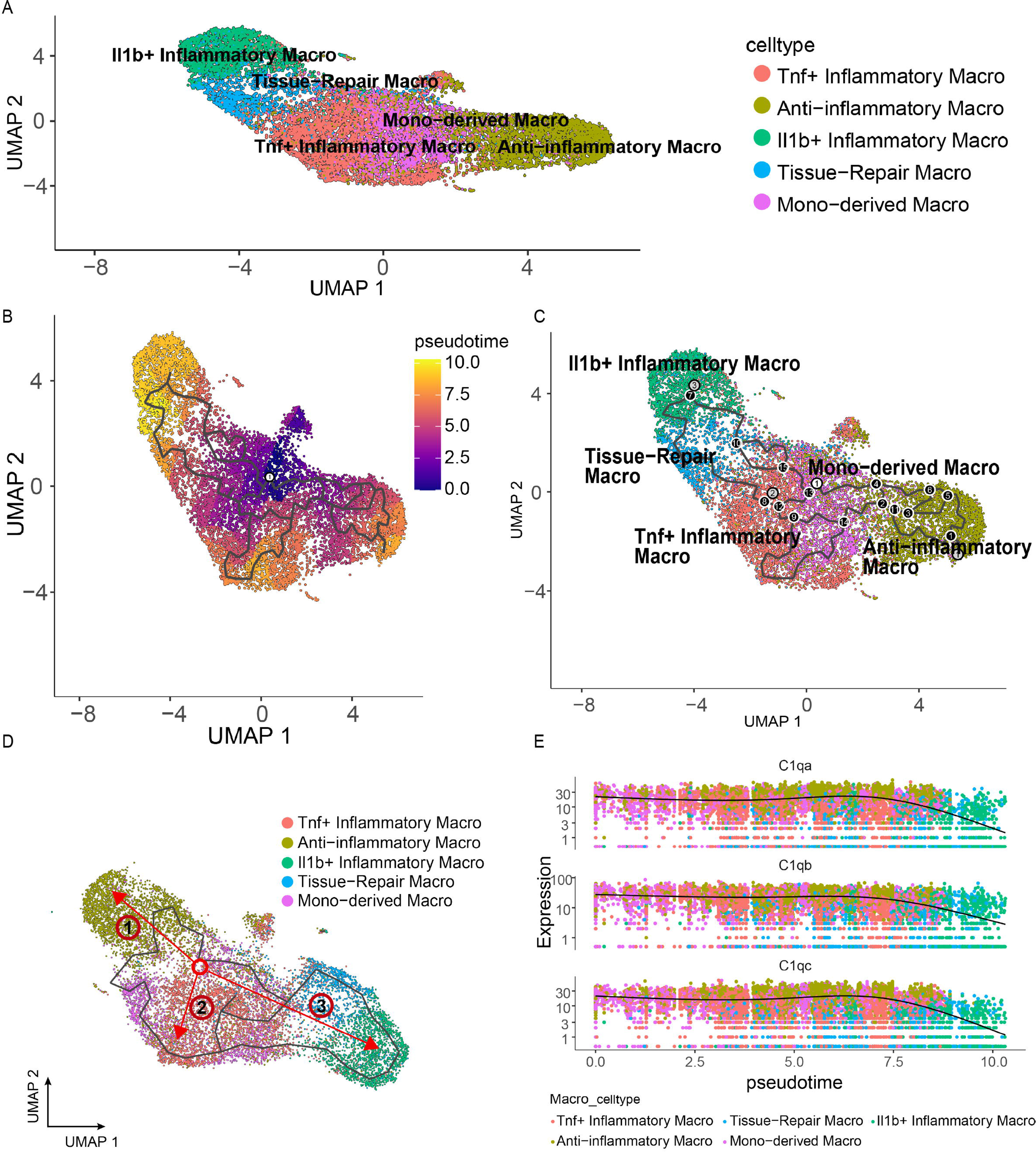
Trajectory analysis of macrophage subtypes. (A) UMAP plot showing 5 macrophage subtypes identified by signature molecular markers; (B) Macrophages’ differentiation trajectories on pseudo-time, with darker purple cells transitioning to lighter orange cells, indicating progression from earlier to later differentiation stages; (C-D) Trajectory analysis revealing dynamic processes across macrophage subpopulations, with RNA expression profiles changing continuously along anatomical axes; numbered circles represent differentiation sequence, where gray-filled circles indicate different endpoints, white-filled circles mark branch points where cells can follow multiple fates, and black-filled numbered circles show different differentiation nodes; (E) Absolute expression of top 3 genes in pseudo-time, where each point represents a cell, different colors represent different clusters, and the y-axis shows gene expression levels.

### 3.6 Cell-to-Cell Communication Between Macrophages and Other Cells

To elucidate macrophage interactions during LA pathogenesis, we analyzed cell-to-cell communication using scRNA-seq data. Macrophage interactions with other cells were significantly enhanced in LA tissue, both in interaction number and strength (Figure 7A-7C). Ligand-receptor binding analysis revealed complex changes: CXCL, VEGF, VISFATIN, MIF, GALECTIN, COMPLEMENT, and OSM showed increased receptor binding, while PTN, CHEMERIN, TNF, IGF, GAS, SEMA3, BAFF, TGF-β, PDGF, CSF, EGF, and CCL exhibited decreased binding (Figure 7D). The heatmap of the TGF-β signaling pathway showed strong associations between anti-inflammatory macro and Il1b+ inflammatory macro and tissue-repair macro (Figure 7E). Network centrality analysis of the inferred TGF-β signaling network identified that all cell populations are sources of TGF-β ligands. Importantly, CellChat predicted that Il1b+ inflammatory macro and tissue-repair macro significantly contributed to TGF-β signal production in LA, revealing a complex TGF-β signaling network with multiple ligand sources targeting a large portion of anti-inflammatory macro. Furthermore, anti-inflammatory macro emerged as the dominant mediator and influencer, suggesting their role as a gatekeeper of cell–cell communication (Figure 7F). The violin plot showed significant expression of the TGFB1 gene across all cell types, and a notable decrease in TGFBR2 expression was observed in anti-inflammatory macro, tissue-repair macro, and Tnf+ inflammatory macro (Figure 7G). Tissue-repair macrophages exhibited high intercellular communication volume in LA (Figure 7H-7I), with the Tgfb1-(Tglbr1+TgIbr2) axis contributing most significantly to TGF-β signaling (Figure 7J). Interestingly, The CellChat showed that the majority of TGF-β interactions among cells are paracrine, with only anti-inflammatory macro, Tnf+ inflammatory macro, and tissue-repair macro demonstrating prominent autocrine signaling (Figures 7K,7L).

**Figure 7.**
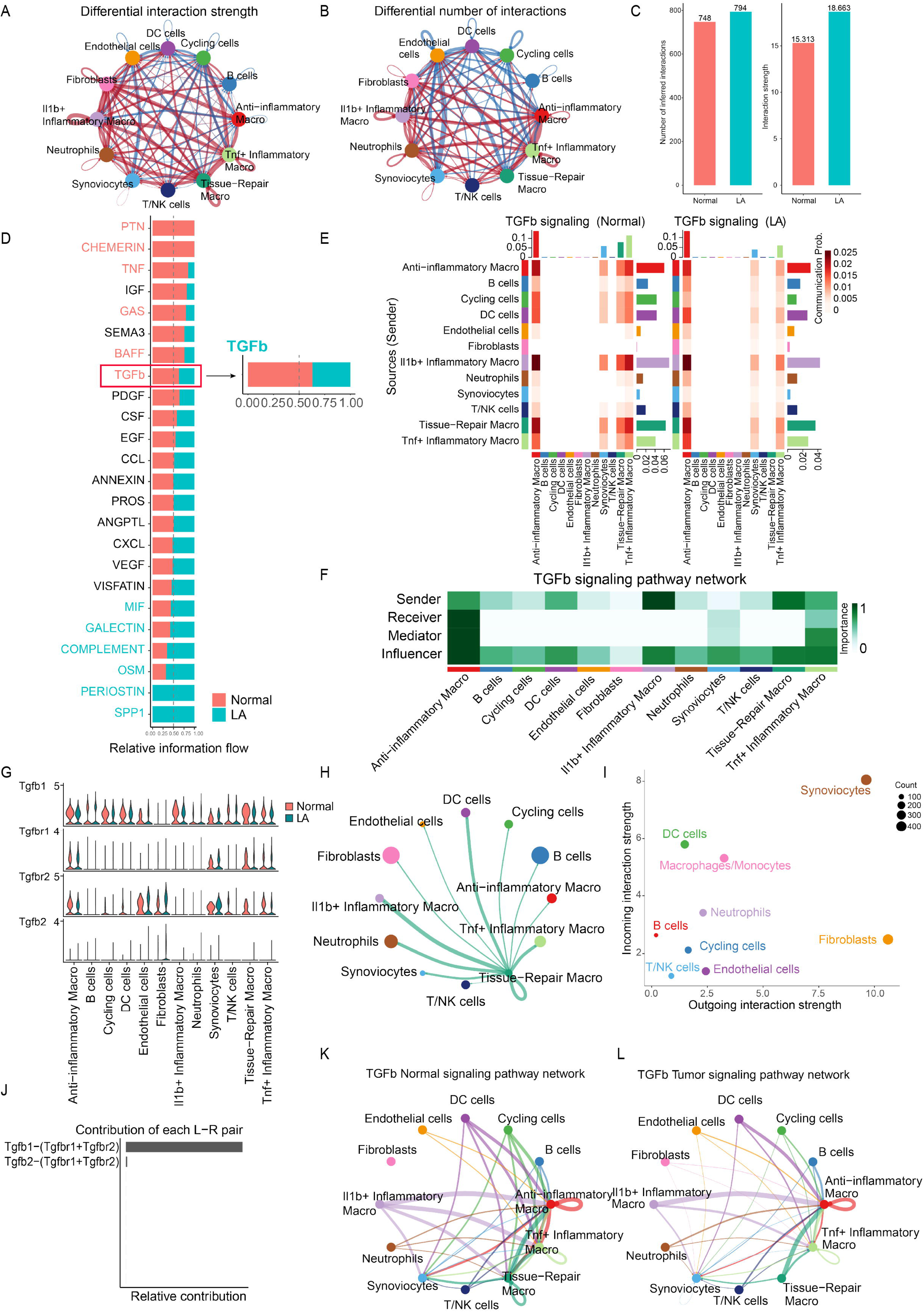
Cellular communication analysis. (A-B) Network diagrams showing cell-cell interaction number and strength in normal and Lyme arthritis model groups respectively; (C) Comparative analysis of cellular communication number and strength between groups; (D) Total pathway analysis comparing Lyme arthritis model and normal groups; (E) Heatmap of ligand-receptor interaction patterns in TGF-β signaling pathway across cell types in both groups; (F) TGF-β signal sending-receiving intensity map in Lyme arthritis group; (G) Expression profiles of representative TGF-β signaling pathway genes across cell types between groups; (H) Interaction network between tissue-repair macrophage and other cell types in Lyme arthritis group; (I) Interaction dynamics among different cell types; (J) Contribution of each TGF-β ligand-receptor pair; (K-L) Cell-cell communication mediated by Tgfb1-(Tglbr1+Tglbr2) ligand-receptor pair in normal and Lyme arthritis model groups.

### 3.7 Bulk-transcriptome sequencing of C57BL/6 Mouse Ankle Joints

To examine macrophage specificity in LA, we merged transcriptome data from C57BL/6 mouse ankle joints with public data from bone marrow-derived macrophages (GSE125503, platform GPL17021) and conducted differential gene analysis. Figure 8A-8B show the heatmap and volcano plot of differential genes. We identified 397 DEGs and intersected them with 222 macrophage-related DEGs from scRNA-seq, yielding 15 overlapping genes (Figure 8C): SIRPB1C, CD14, FABP5, IL2RG, MCU, MMP14, MXRA8, EGR1, MSR1, CLEC4D, PIRB, IL1B, H2-AA, GYPC, and ITGA6. After normalizing the expression matrix, LASSO regression analysis identified four candidate optimal feature genes: SIRPB1C, FABP5, MMP14, and EGR1 (Figure 8D-8E), with their expression profiles shown in Figures 8F-G-H-I.

**Figure 8.**
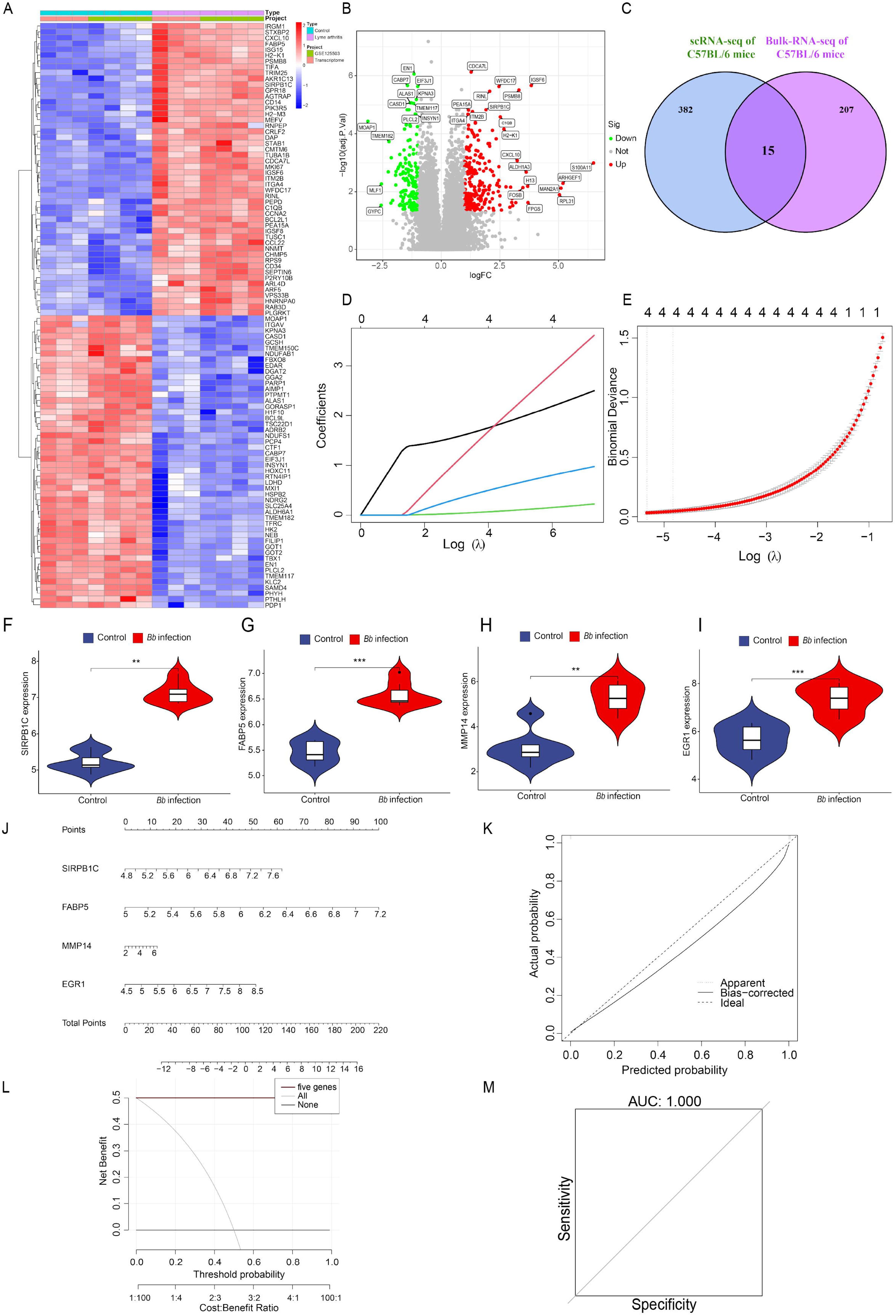
Machine learning prediction of Lyme arthritis candidate optimal feature genes and nomogram for disease occurrence prediction. (A) Heatmap of differentially expressed genes in merged transcriptome: blue means low expression; red means high expression; (B) Volcano plot of differentially expressed genes in merged transcriptome: green means low expression; red means high expression; (C) Venn diagram identified the 15 overlapping genes between merged transcriptome DEGs and Macrophage/Monocyte-related genes from single-cell data; (D) LASSO regression model construction using merged transcriptome samples as training set; (E) Model parameter optimization cross-validation (left line indicating optimal λ value), determined the four candidate optimal feature genes (SIRPB1C, FABP5, MMP14 and EGR1); (F-J-H-I) Expression levels of the 4 candidate optimal feature genes in merged transcriptome samples; (J) Nomogram for Lyme arthritis occurrence prediction. Line segments represent the predicted contribution of genes to outcome events, with the total score signifying the cumulative sum of all corresponding individual scores across variable values; (K) Calibration curve evaluating nomogram model predictive ability; (L) DCA curve assessing nomogram model value; (M) ROC curve evaluating nomogram model performance in merged transcriptome samples. \**p* < 0.05, \*\**p* < 0.01, \*\*\**p* < 0.001.

### 3.8 Construction and testing of a signature gene-based nomogram for predicting LA

Using the “RMS” R package, we constructed a diagnostic nomogram for LA based on the four characteristic genes (Figure 8J) and evaluated its predictive capability. The calibration curve showed minimal difference between actual and predicted LA risk, indicating high model accuracy (Figure 8K). Decision curve analysis (DCA) demonstrated potential patient benefit from this nomogram (Figure 8L), while ROC curve analysis further confirmed the model’s accuracy (Figure 8M).

### 3.9 Validation of Candidate Optimal Feature Genes

In our LA mouse model, C57BL/6 mice developed hind leg ankle joint swelling 2 days after Bb injection, peaking at approximately 2 weeks before gradually subsiding after 4-5 weeks. H&E staining showed inflammatory cell infiltration and substantial edema in joint synovial and surrounding soft tissues, while bone and cartilage remained intact (Figure 9A). By day 16, experimental mice displayed significant joint swelling and mild stiffness compared to PBS controls, without skin ulceration or blackening (Figure 9B). X-rays revealed soft tissue edema (Figure 9C). RT-qPCR analysis confirmed significantly increased expression of SIRPB1C, FABP5, MMP14, and EGR1 in Bb-infected samples from both RAW264.7 cells and joints of C57BL/6 mice compared to controls.

**Figure 9.**
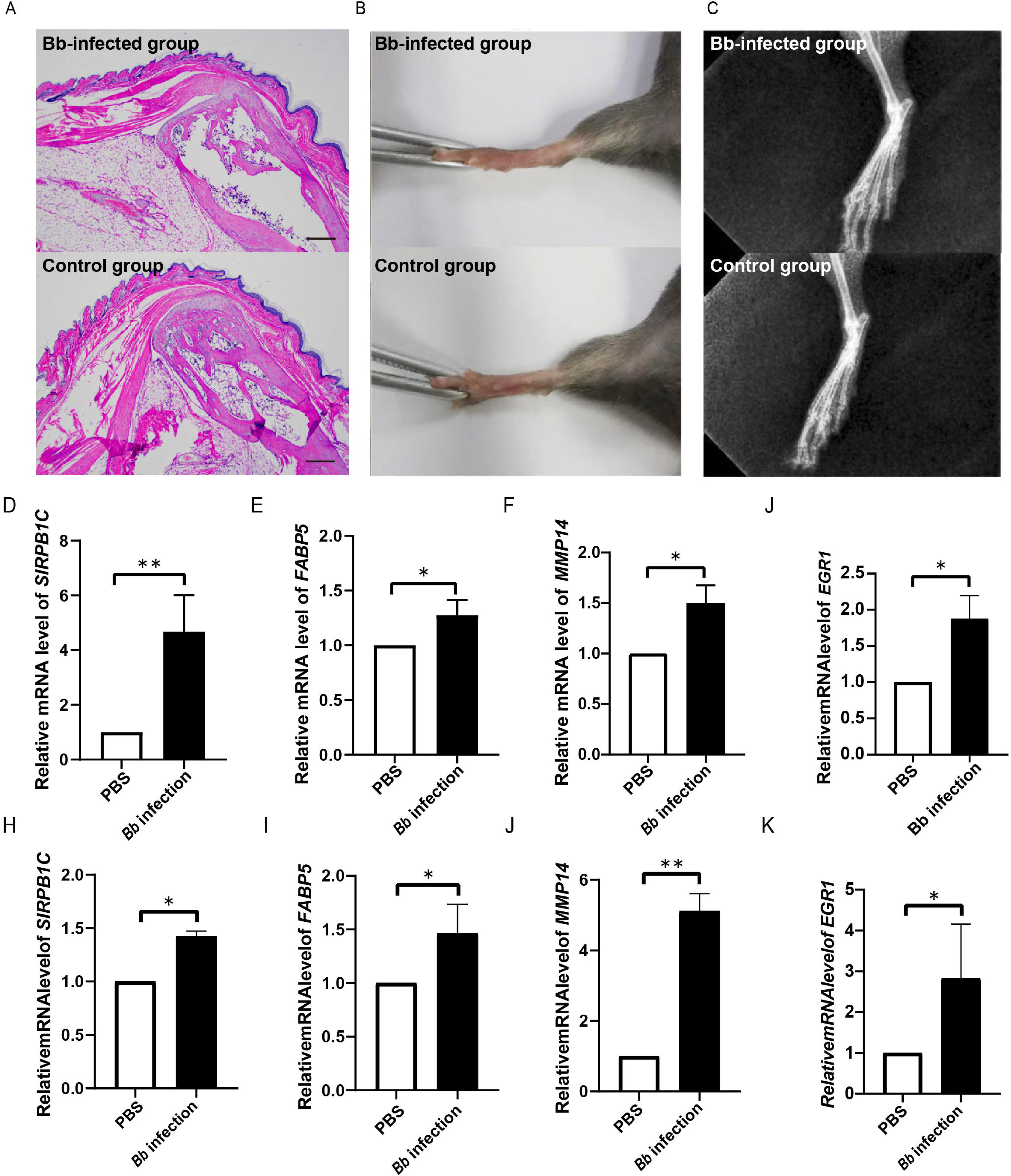
Validation of 4 candidate optimal feature genes. (A-B-C) Edema observation, X-ray examination, and H&E staining of joint synovium in Lyme arthritis mouse model versus control, scale bar = 750 μm; (D-K) RT-qPCR validation of these genes in Borrelia burgdorferi–infected RAW264.7 cells and C57BL/6 mice versus controls. *p < 0.05, **p < 0.01, ***p < 0.001.

## 4. Discussion

LA is one of the most common late-stage manifestations of Lyme disease. Without timely diagnosis and treatment, it can lead to persistent arthritis and long-term functional impairment [33]. While antibiotics are generally effective, 10-20% of patients develop antibiotic-refractory LA (ARLA) [34,35]. Single-cell RNA sequencing (scRNA-seq) now enables analysis of disease pathogenesis at unprecedented resolution [36,37], revealing cellular composition and transcriptomic changes associated with LA. Macrophages, as key effector cells of the innate immune system, play a central role in the pathogenesis of LA [38], regulating inflammatory responses and influencing disease progression through cytokine and chemokine secretion. C57BL/6 mice provide an ideal model for studying transcriptional and cellular changes during infection due to their uniform genetic background. In this model, arthritis symptoms resolve spontaneously despite persistent bacteria, effectively simulating host immune adaptation to chronic infection[39–42]. As the disease progresses, bacterial numbers gradually decrease while the host immune response changes, achieving long-term control of infection[18]. This self-healing process provides valuable insights into the self-limiting nature of LA. This study analyzed macrophage differential gene expression in LA using GSE233850 scRNA-seq data from C57BL/6 mouse models. By integrating bulk transcriptome datasets through joint analysis and machine learning, we identified and validated core macrophage-related genes in LA. The synovial microenvironment (SME), comprising immune cells, synovial cells, and stromal cells[43], critically influences LA progression and prognosis[44]. Antibiotic-refractory LA patients often experience poor outcomes[5], with some showing persistent macrophage activation despite Bb clearance[45]. Since synovial macrophages can either promote or inhibit inflammation through M1/M2 polarization, understanding these mechanisms is essential for improving treatment[46]. Single-cell RNA sequencing has become vital for characterizing synovial immune cells in inflammatory arthritis[47,48]. Therefore, systematically studying macrophage functional states and regulatory networks in LA, and developing macrophage-based prognostic models, has significant clinical relevance.

Single-cell transcriptome analysis of ankle joints from uninfected and Bb-infected C57BL/6 mice identified 9 cell types, with significant shifts during LA development: Fibroblasts increased while Macrophages/Monocytes decreased. We found seven Macrophage/Monocyte subsets with notable compositional differences between normal and LA models. LA samples showed decreased anti-inflammatory macrophages, monocyte-derived macrophages, and Ly6c+ inflammatory monocytes, with increased TNF+ and Il1b+ inflammatory macrophages, tissue-repair macrophages, and progenitor cells. These subsets serve distinct functions: anti-inflammatory macrophages maintain homeostasis by suppressing inflammation[49–51]; TNF+ inflammatory macrophages mediate inflammatory and chemokine signaling[52–54]; tissue-repair macrophages orchestrate inflammation regulation and tissue remodeling[55,56]. Monocyte-derived macrophages play a crucial regulatory role in various diseases, particularly in liver regeneration, where they facilitate tissue repair through migration, proliferation, and apoptosis[57].In neuroinflammation, such as stroke, these cells migrate to the brain via CCR4 and CCR5 receptors, emerging as potential therapeutic targets[58]; IL-1β+ inflammatory macrophages initiate inflammatory cascade [59–61]; Ly6c+ inflammatory monocytes drive inflammation and angiogenesis[62,63]; and monocytes/progenitor cells respond to systemic inflammation in various diseases[64–66]. Despite the overall reduction in Macrophages/Monocytes under pathological conditions, IL1b+ inflammatory and tissue-regenerative macrophages significantly increased while anti-inflammatory macrophages declined, indicating functional reprogramming within the macrophage spectrum. Despite overall Macrophage/Monocyte reduction, IL1b+ inflammatory and tissue-regenerative macrophages increased while anti-inflammatory macrophages declined, indicating functional reprogramming. Pseudo-temporal trajectory analysis revealed three main differentiation pathways from monocyte-derived macrophages: anti-inflammatory, TNF+ inflammatory, and IL1b+ inflammatory lineages.

Cell-cell communication analysis identified significant changes in Transforming growth factor-β (TGF-β)-receptor interactions during pathological states, suggesting potential therapeutic targets for LA. Heat map analyses showed decreased expression of TGF-β signaling pathway-related genes between various cell types in LA compared to normal tissues. We identified the critical ligand-receptor pair Tgfb1-(Tglbr1+TgIbr2) within TGF-β signaling networks showing enhanced intercellular communication across diverse cell types in LA. TGF-β, a cytokine that regulates inflammation and promotes fibrosis[67], plays a dual role in Lyme disease. TGF-β1 levels are higher in early/non-chronic patients compared to chronic patients, who show impaired TGF-β1 synthesis when stimulated by spirochetes[68]. This difference may explain the mechanism of disease progression to chronicity: elevated TGF-β1 levels represent an effective anti-inflammatory immune response, promoting pathogen clearance; whereas the reduction of TGF-β1 in chronic patients leads to insufficient persistent anti-inflammatory response[69]. Patients with antibiotic-refractory LA show excessive inflammation and fibrosis in synovial tissue[70]. TGF-β affects vascular reconstruction and may promote vascular occlusion[71], suggesting occlusive microvascular lesions in antibiotic-refractory LA patients may be linked to TGF-β activity.

GSVA and KEGG pathway analyses showed that Bb infection shifted macrophage metabolism from oxidative phosphorylation to glycolysis. Previous studies confirm metabolic reconfiguration’s critical role in inflammation: lipopolysaccharide-induced metabolic transition is essential for inflammatory response[72]; while increased glycolysis enhances macrophage inflammatory functions in ankylosing spondylitis and myocardial infarction[73] [74]. These findings support our conclusions about metabolic reprogramming of Monocytes/Macrophages in LA.

We developed a LA model in C57BL/6 mice and analyzed transcriptome data from infected ankle joints combined with public dataset GSE125503 to investigate macrophage roles in LA. Through differential gene expression and LASSO regression, we identified four characteristic genes: SIRPB1C, FABP5, MMP14, and EGR1. (Signal Regulatory Protein Beta 1C (SIRPB1C) belongs to the signal regulatory protein family involved in inflammation and immunity. SIRPB1C is a key member of the signal regulatory protein beta family. Studies show SIRPB1 knockout reduces inflammatory cytokines (IL1RA, CCL2, IL-8) [75], while another family member, SIRPα, suppresses phagocytosis and immune cell migration, with SIRPα-targeted antibodies reducing inflammation[76,77]. Fatty acid-binding protein 5 (FABP5), a lipid chaperone involved in inflammatory pain modulation, is present in human synovium[78,79]. Its inhibition significantly reduces cytokine and chemokine secretion, suggesting potential impact on osteoarthritis-related inflammatory pain[80]. FABP5 is upregulated in intestinal macrophages during inflammatory bowel diseases, and its inhibition prevents inflammatory M1 macrophage differentiation, reducing colitis in mice[81].

MMP14 (membrane type I matrix metalloproteinase or MT1-MMP) is a collagenase that degrades extracellular matrix, particularly type II collagen in cartilage[82]. In rheumatoid arthritis, MMP14 on synoviocytes and macrophages promotes cartilage degradation[83]. Studies show MMP14 inhibition preserves joints: the inhibitor FR255031 reduces cartilage degradation in arthritis models, while selective MT1-MMP inhibition protects joints and enhances anti-tumor necrosis factor therapy efficacy, suggesting potential combination treatments[84,85].

Early Growth Response 1 (EGR1), a transcription factor, plays a crucial role in host-pathogen interactions[86]. While no direct EGR1-LA link has been previously documented, this factor is rapidly induced by pathogens, cytokines, and stressors, and regulates inflammation, angiogenesis, and matrix remodeling[87,88]. EGR1 promotes cartilage angiogenesis through the NT-1/DCC-VEGF pathway, accelerating cartilage deterioration[89]. As an inflammatory mediator, it enhances pro-inflammatory IL-1 effects while suppressing anti-inflammatory PPARγ[90]. EGR1 also regulates extracellular matrix genes, affecting cartilage integrity[91]. Our research suggests EGR1 likely plays an essential role in LA inflammatory regulation.

Although our research provides valuable insights, it has several limitations that need to be addressed in future studies. First, the sample size of the integrated bulk RNA sequencing data is relatively limited. Second, regarding the exploration of molecular mechanisms: while we validated the transcriptional changes of core differential genes through in vitro and vivo experiments, we have not fully understood the functional roles of these genes in LA pathogenesis. Future research should aim to validate these findings in larger cohorts and employ functional experiments to confirm the roles of these cells, genes, and pathways in disease progression.

## 5. Conclusion

Integrative single-cell and bulk RNA sequencing revealed seven macrophage subtypes in LA, a glycolytic shift, and disrupted TGF-β signaling. Pseudo-time analysis traced monocyte-derived macrophages toward distinct anti-inflammatory or TNF+/IL1β+ inflammatory fates, underscoring functional plasticity. Four candidate optimal feature genes-SIRPB1C, FABP5, MMP14 and EGR1-accurately discriminated diseased joints, forming a nomogram with high predictive value. These findings deepen understanding of macrophage heterogeneity in LA and highlight metabolic and signaling nodes that may guide biomarker-driven diagnosis and macrophage-targeted therapy.

## Acknowledgement

We thank Professor Yi[Qun Kuang for providing the Biosafety Laboratory at the Research Center for Clinical Medicine, First Affiliated Hospital of Kunming Medical University, and acknowledge the GEO databases for publicly available datasets.

## Contributors

Yan Dong and Yantong Chen performed bioinformatics analyses and drafted the manuscript; Yanshuang Luo, Meng Liu, Yingchen Zhou, Yutian Yang, Xuesong Chen, Xiaorong Liu, Fusong Yang, Zhi Liang, Liye Zi, Hongbing Gao, Zhichao Cao, and Fucun Gao conducted experiments, interpreted data; Guozhong Zhou and Chao Song designed the study and supervised the project. All authors reviewed and approved the final manuscript.

## Funding

This study was supported by the Yunnan Fundamental Research Projects (grant NO. 202301BE070001-036) and internal projects of the First People’s Hospital of Anning, affiliated with Kunming University of Science and Technology (grant NO. 2024AYY001 and 2024AYY005).

## Competing interests

None declared.

## Ethical approval

Animal experiments were conducted in accordance with the Declaration of Helsinki principles and approved by the Kunming Medical University Experimental Animal Ethics Committee (Approval No.: KMMU20211590).

## Provenance and peer review

Not commissioned; externally peer reviewed.

## Data availability statement

Publicly available datasets that support the findings of this study are openly available in the Gene Expression Omnibus (GEO) database (https://www.ncbi.nlm.nih.gov/geo/info/datasets.html); GSE233850 and GSE125503.

## Code availability

Additional information for data re-analysis is available from the lead contact upon request.

## Publisher’s note

Claims expressed in this article are solely those of the authors and do not necessarily represent affiliated organizations or the publisher. Products evaluated or claims made by manufacturers are not guaranteed or endorsed by the publisher.

## Notes

### Competing Interest Statement

The authors have declared no competing interest.

### Summary of Updates

Introduction Lyme arthritis (LA) is a common late-stage manifestation of Lyme disease caused by Borrelia burgdorferi (Bb) infection. Macrophages play a crucial role in LA pathogenesis, yet their heterogeneity and functional states within the arthritic microenvironment remain poorly understood. Methods We integrated single-cell RNA sequencing data from ankle joint tissues of Bb-infected C57BL/6 mice with bulk RNA sequencing data from both infected ankle joints and bone marrow-derived macrophages. We developed a LA mouse model and validated key findings through histopathological examination and RT-qPCR analysis. Results We identified nine distinct cell types in the joint microenvironment with significant compositional shifts during disease progression. Analysis of 16,617 Macrophages/Monocytes revealed seven functionally distinct subpopulations with differential responses to infection. Pseudo-time trajectory analysis demonstrated three principal differentiation pathways from monocyte-derived macrophages toward anti-inflammatory, TNF+, and IL1b+ inflammatory phenotypes. Cell-cell communication analysis revealed significantly altered TGF-β signaling networks in LA, with the Tgfb1-(Tglbr1+TgIbr2) axis playing a critical role. GSVA and pathway analyses showed metabolic reprogramming from oxidative phosphorylation toward glycolysis in macrophages during infection. Through integrated analysis and LASSO regression, we identified four characteristic genes (SIRPB1C, FABP5, MMP14, and EGR1) as potential LA biomarkers and developed a diagnostic nomogram with high predictive accuracy. Conclusion Our integrative analysis provides comprehensive insights into macrophage heterogeneity and functional plasticity in LA, identifying characteristic biomarkers and potential therapeutic targets. The metabolic reprogramming and altered TGF-β signaling networks we identified may contribute to disease pathogenesis and offer new avenues for intervention strategies.

